# Retrograde Optogenetics Reveals Functional Convergence within a Corticotectal Pathway of Non-Human Primates

**DOI:** 10.1101/2025.08.03.667949

**Authors:** Xuefei Yu, Atul Gopal, Ken-ichi Inoue, Martin Bohlen, Genevieve Kuczewski, Marc A. Sommer, Hendrikje Nienborg, Masahiko Takada, Okihide Hikosaka

## Abstract

Understanding how the cerebral cortex communicates with subcortical areas to drive behavior remains a central question in system neuroscience. One key unresolved issue is whether prefrontal cortical outputs to motor-related subcortical regions carry predominantly motor commands or mixed sensory-motor signals. Retrograde optogenetics offers a powerful way to interrogate such projection-defined circuits, but its use in non-human primates has been limited. Here, we applied retrograde optogenetics in awake macaques to directly test the functional organization of the corticotectal projection from the frontal eye field (FEF) to the superior colliculus (SC). We asked whether the FEF output signals to SC are motor-dominant or broadly sensory-motor. Optical activation of this pathway evoked robust, contralateral saccades and selectively modulated reaction times, demonstrating its causal role in saccade generation. Optogenetically tagging FEF neurons projecting to SC revealed a heterogeneous population of visual, visuomotor, and motor neurons. This diverse output converged predominantly onto motor-related neurons in the SC. These findings support a visuomotor convergence model, in which diverse FEF outputs drive motor-selective SC neurons with activity sufficient for saccade generation, and thus resolve long-standing questions over the composition of FEF outputs. Additionally, our results establish retrograde optogenetics as a tool for dissecting projection-defined circuits in primates and for precisely probing the neural pathways that link perception to action.

**Highlights:** *Retrograde optogenetics enabled pronounced and selective modulation of behavior in macaques*.

*Selective optical stimulation of the FEF–SC pathway evoked contralateral saccades and modulated reaction times*.

*Opto-tagging revealed diverse visual, visuomotor, and motor FEF outputs converging onto motor-related SC neurons*.

*Findings support a visuomotor convergence model and demonstrate the power of projection-specific optogenetic tools in primates*.

## Introduction

What information does the cerebral cortex transmit to subcortical structures to guide actions? This fundamental question is central to understanding how distributed and complex brain networks convert perception and cognition into action. In non-human primates, the prefrontal cortex is often regarded as a core region where diverse sensory and contextual information is flexibly integrated to guide decisions and actions^1^. However, whether its output to subcortical action-oriented areas primarily conveys the *results* of this integration, i.e. motor-dominant signals, or instead an *intermediate stage*, i.e. sensory-motor signals, remains debated.

One example from everyday life is visual inspection of a scene and the decision where to look next. This process is thought to rely largely on output from the frontal eye field (FEF), a prefrontal region critical in cognitive control^2^, to the superior colliculus (SC), a brainstem structure essential for generation of eye movement^3^. Early studies portrayed this projection as carrying predominantly motor commands^4^, whereas subsequent research indicated a richer signal composition, encompassing visual, visuomotor, and motor signals^5,6^. The functional makeup of the FEF–SC projection therefore remains controversial. Moreover, FEF-SC signals have never been directly characterized in parallel with causal tests of their role on behavior.

Here we applied retrograde optogenetics^7-10^ in rhesus macaques^11-14^ to selectively stimulate the FEF-SC pathway and determine its influence on saccade generation. Activation of the SC-projecting FEF neurons strongly modulated saccades. Recordings from the optically modulated neurons in FEF revealed that its SC-projecting neurons convey diverse visual, visuomotor, and motor signals that converge onto motor-related SC neurons. Our findings support a model of visuomotor convergence in the FEF–SC pathway and establish a generalizable framework for applying retrograde optogenetics in the primate brain to understand perception-action circuits.

## Results

### Pronounced and selective behavioral modulation by retrograde optogenetics

We injected AAV2retro-hSyn-ChR2-EYFP unilaterally into the SC of two monkeys (Figure 1A). To confirm gene expression in the FEF-SC projection, we successfully manipulated neural activity and saccadic behavior using fiber optic illumination of the SC (Figure S1A-H) and showed post-mortem expression of the ChR2-EYFP complex via histological analysis of the SC (Figure 2A) and FEF. Expression was profuse in the ipsilateral FEF (Figure 2B) but negligible in the contralateral FEF (Figure 2C).

**Figure 1.**
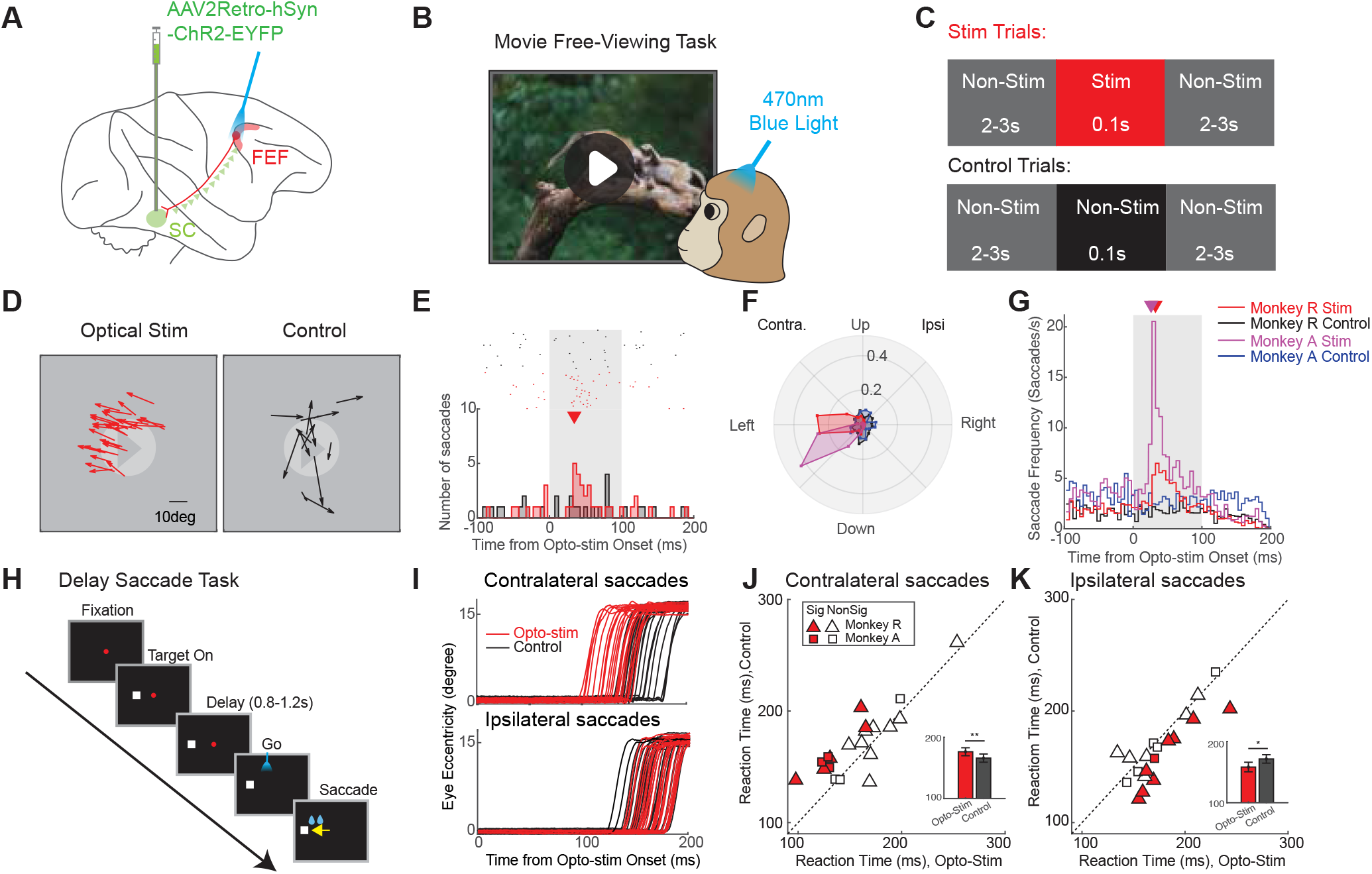
Robust saccade modulation induced by optical stimulation of the FEF-SC pathway. **(A)** The retrograde viral vector was injected into the SC, from where it was transported retrogradely to the somata of SC-projecting neurons in the FEF. Optical stimulation was applied in the FEF. **(B)** Free-viewing task: macaques viewed movies and were rewarded for keeping their gaze within the display. **(C)** Optical stimulation protocol during free-viewing. Stimulation trials (top) included a 0.1s optical pulse from a blue laser. Control trials (bottom) included a 0.1s non-stimulated period. Both were flanked by 2-3s non-stimulated periods. **(D)** Example optical stimulation session in FEF showing saccades that occurred in stimulation (left) and control (right) trials. **(E)** Example peri-stimulus time histogram (PSTH) of saccades in the stimulation (red) and control trials (black) during optical stimulation in the FEF (N = 40 trials per condition). Upper raster shows time of saccade onsets in individual trials (both spontaneous and putatively stimulation-evoked). Lower histogram shows distributions of those onset times binned at a 5ms intervals. Triangle is time of peak saccade occurrence (35ms). **(F)** Polar histograms of saccade directions in stimulation and control trials for both monkeys during optical stimulation in the FEF. The radial (ρ) axis represents the proportion of saccades within each angular bin (width: 22.5°), normalized to the total number of saccades recorded during the 100ms interval under each condition for each monkey. **(G)** Population PSTH of saccades in stimulation and control trials for both monkeys during optical stimulation in the FEF. To generate the PSTH, saccades were first counted and averaged across trials in 5ms bins from -100ms to 200ms relative to stimulation onset for each session. The average counts were then divided by the 5ms bin duration to yield the saccade frequency. The triangle indicates the time of peak saccade occurrence for monkey R (red: 32.5ms) and monkey A (magenta: 27.5ms). **(H)** Delayed saccade task. In each trial, a target appeared in the left or right visual field, followed by a delay period of 0.8-1.2s. Then the fixation point disappeared, cueing the animal to make a saccade to the target location. **(I)** Example session showing eye movement traces for contralateral (top) and ipsilateral saccades (bottom) during optical stimulation (red) and control (black) trials. Reaction times: Contralateral, 122.9 ± 8.1ms [Mean ± S.E.M] for Stim vs. 154.4 ± 3.3ms for Control, p = 7.45e-04; Ipsilateral, 170.4 ± 5.7ms for Stim vs. 157.6 ± 5.7ms for Control, p = 0.06; all N = 20 trials. **(J)** Scatter plot of mean reaction times across sessions (N = 20) for contralateral saccades in optogenetic Stim vs. Control trials. Filled Symbols: Individual sessions with a significant difference between Stim and Control (p < 0.05; Triangles: Monkey R; Squares: Monkey A. Inset: pooled comparisons [Mean ± S.E.M]. At this population level, too, reaction times were shorter for Stim (158.23 ± 7.81ms) than for Control (171.44 ± 6.89ms; p = 0.0068). **(K)** Same but for ipsilateral saccades, which in both individual sessions and in the pooled data showed longer reaction times in Stim vs. Control trials. For inset, pooled data: Stim 176.43 ± 6.54ms vs. Control 166.04 ± 6.69ms; p = 0.0084.

**Figure 2.**
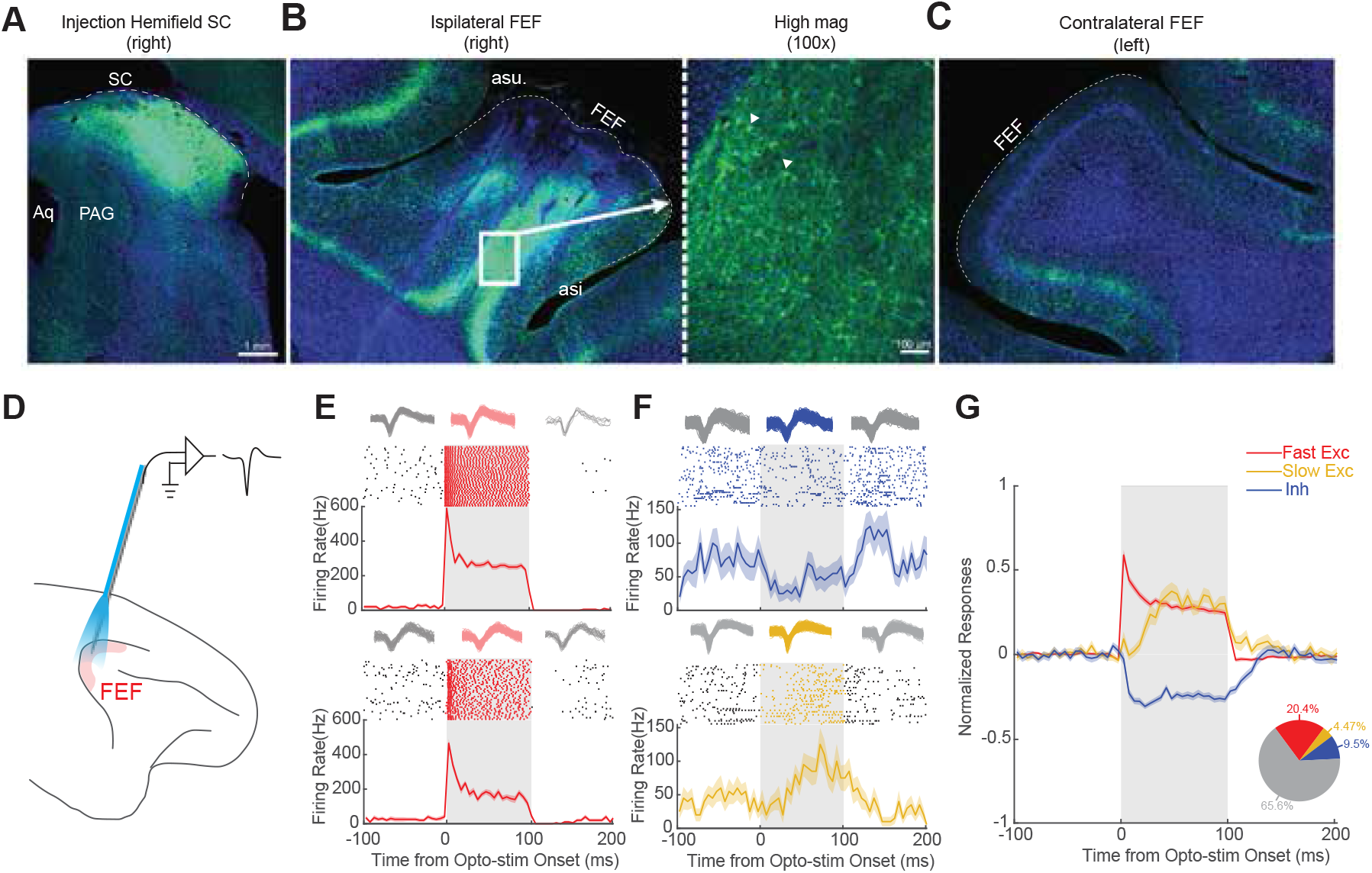
AAV2retro efficiently labeled SC-projecting FEF neurons. **(A)** Immunofluorescence showing ChR2-EYFP expression in the injected hemifield of SC. **(B)** Immunofluorescence showing retrograde labeling of ipsilateral FEF neurons by ChR2-EYFP. The right panel is the magnified window(100X) showing the labeled somata. The triangles indicate two examples. **(C)** Negligible immunofluorescence in the contralateral FEF. **(D)** A schematic illustrating simultaneous optical stimulation and neuron recording in the FEF. **(E)** Two examples of putative direct activation by optical stimulation (“Fast Exc” neurons). Top: spike waveforms, collected during the 100ms optical stimulation period (red) or outside it (gray), demonstrating stable unit identity. Middle: rasters of spike times. Bottom: spike density functions (SDF). **(F)** Same but neurons showing putative indirect activation. Top neuron: Inhibition (“Inh”). Bottom neuron: Delayed activation (“Slow Exc”). **(G)** Population average across the Fast Exc, Slow Exc, and Inh neurons in the FEF. The raw spike density function (SDF) for each neuron was normalized by dividing by its peak firing rate and then the average baseline activity (-100ms to 0ms relative to laser onset) was subtracted. Inset: Pie chart showing the proportion of FEF neurons in each group. Fast Exc (red), N = 146; Slow Exc (orange), N = 32; Inh (blue), N = 68; No effect (gray), N = 470.

Our first goal was to optically stimulate the FEF to test whether selective activation of SC-projecting neurons evoked saccades. Monkeys freely viewed a movie (Figure 1B) while light (0.1s, 470nm, 50mW/mm^2^) was delivered intermittently to the FEF (Figure 1B,C). Saccades were compared between these “Stim Trials” (Figure 1C, top) and non-stimulated “Control Trials” (Figure 1C, bottom). Across 110 sessions, saccade frequency significantly increased during the optical stimulation period (4.24 saccades/s, N = 2443 saccades) relative to the control period (2.41 saccades/s, N = 1408 saccades, p = 4.28e-11). Evoked saccades were consistently directed toward the left visual field (example session in Figure 1D), contralateral to the site of stimulation (see also Figure S2A), similar to electrically stimulated saccades from the same sites (Figure S2B-D). The onset latency of these example optically evoked saccades from FEF was 35ms post-stimulation (Figure 1E), about 12ms longer than the latency observed from optical stimulation in SC in the same monkey (22.5ms, Figure S1F,H). Saccadic vectors evoked from optical stimulation in FEF and the SC were generally well aligned (Figure 1F compared with Figure S1G). The FEF optical stimulation results were highly reproducible across sessions and animals (Figure 1F, Figure S2A), with a consistent latency peak ranging from 27.5 ms to 32.5ms following laser onset (Figure 1G), compared with 22.5ms latency from SC (Figure S1H). This 5 - 10ms latency difference is consistent with axonal conduction and synaptic delay through the FEF-SC pathway^15,16^. Optical stimulation of the FEF in fixation tasks also induced consistent contralateral saccades, though with smaller amplitudes and reduced frequency (Figure S2E-J). Importantly, no such effects were observed prior to the viral vector injection (Figure S2K) or during red light stimulation, ineffective to activate ChR2 (Figure S2L), confirming the specificity of the ChR2-mediated effects.

Next, we investigated whether subthreshold optical stimulation in the FEF could modulate the planning of saccades without directly evoking them, as found in electrical stimulation studies of the FEF and SC^17,18^. We applied low-intensity optical stimulation (< 20mW/mm^2^) during a delayed saccade task (Figure 1H) targeting the final phase of saccade preparation (200ms after fixation offset). For contralateral saccades, reaction times decreased with stimulation (Stim 158.23 ± 7.81ms [Mean ± S.E.M] vs. Control 171.44 ± 6.89ms, both N = 20 sessions, p = 0.0068, Figure 1I,J). The facilitation effect was strongest when the target location was aligned with the direction of the evoked saccade vector (Figure S2M). By contrast, ipsilateral saccades were delayed (Stim 176.43 ± 6.54ms vs. Control 166.04 ± 6.69ms, both N = 20 sessions, p = 0.0084; Figure 1I,K). This effect was most pronounced when the target was positioned opposite to the sites’ preferred direction (Figure S2M). During this manipulation, optical stimulation did not evoke additional saccades, confirming that the stimulation remained subthreshold and modulated the reaction time in a selective way.

Taken together, these findings demonstrated the efficacy of our approach and confirmed the sufficiency of the FEF-SC pathway for generating saccades (Figure 1D-G) and modulating their timing (Figure 1H-K). Next, we leveraged opto-tagging to characterize the signals in the pathway.

### Identification of FEF neurons that project to the SC

Our retrograde optogenetic approach provides a new way to evaluate two competing hypotheses about cerebral cortical output to subcortical areas, using the FEF projection to SC as a test case. Are the output signals preferentially motor-related,^4^ or do they span the full visuomotor continuum^5,6^? To test this, we conducted simultaneous optical stimulation and electrophysiological recording in the FEF using a customized optrode^19^ (Figure 2D). A substantial proportion of FEF neurons (34.4%, 246/716 from both monkeys) showed robust modulation in response to optical stimulation (examples in Figure 2E,F), most commonly with increased firing rates. A subset of neurons (20.4%, 146/716) exhibited rapid, pronounced activation at <10ms latency following laser onset (Figure 2E,G; Figure S3A). We considered these to be SC-projecting FEF neurons directly activated via ChR2 (“Fast Exc”; note, a <5ms latency cutoff yielded similar results; Figure S3G left). We also observed neurons exhibiting either inhibition (“Inh”; Figure 2F, upper panel) or delayed activation (“Slow Exc”; Figure 2F, lower panel) consistent with indirect modulation via synapses^20,21^. The presumed directly and indirectly modulated neurons also showed different strengths of optical response modulation (Figure S3B, Modulation strengths: Fast Exc, 65.54 ± 7.14, N = 146; Slow Exc, 16.32 ± 3.03, N = 32; Inh, 12.44 ± 0.87, N = 68; p = 7.84e-36). None of these effects were observed prior to the viral vector injection (Figure S3C,E) or during red-light stimulation which is ineffective for ChR2 activation (Figure S3D,F), ruling out nonspecific thermal effects. Subsequent analyses focused on the “Fast Exc” category of putatively SC-projecting FEF neurons.

### Signal content of FEF neurons that project to the SC

The FEF-SC pathway can be conceptualized in three alternative organizational schemes, based on the functional properties of FEF output neurons and the properties of FEF-recipient SC neurons (Figure 3A). At one extreme, the *motor dominance model* posits that FEF outputs are primarily motor-related signals^4^ that target visuo-movement and movement neurons in the SC (Figure 3A, left). At the other extreme, the *no convergence model* proposes that the FEF transmits the full spectrum of signals, from purely visual to purely motor^5,6^, and broadly targets all SC neuron types (Figure 3A, right). An intermediate, *visuomotor convergence model* also proposes full-spectrum output from the FEF, but which converges onto movement-related neurons in the SC (Figure 3A, middle). To distinguish between these models, we characterized both the functional properties of SC-projecting FEF neurons and their targets within the SC.

**Figure 3.**
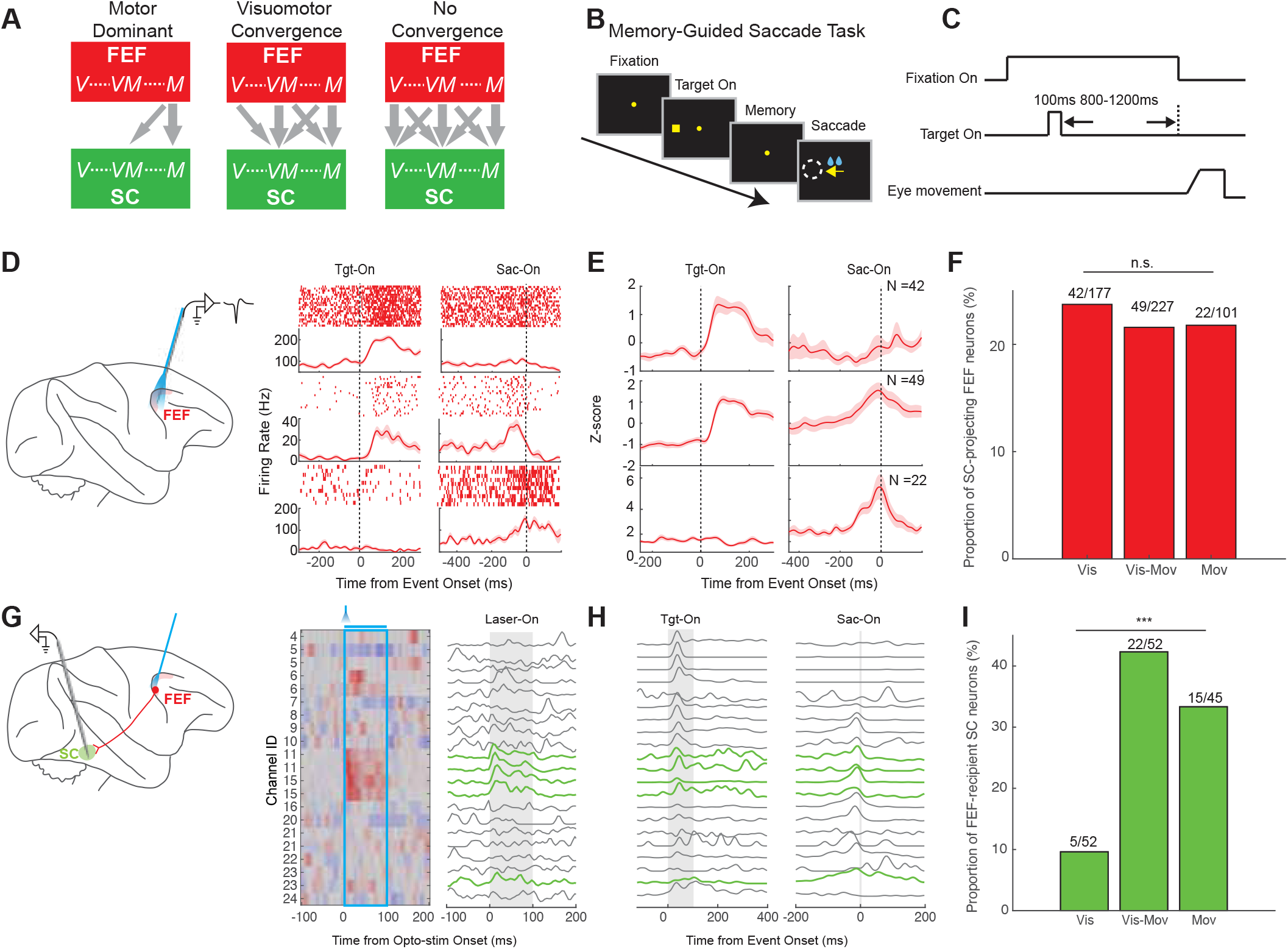
Signal transformations in the FEF-SC circuit for saccade generation. **(A)** Schematic illustrating possible models of information flow from the FEF to the SC. Both the FEF and SC contain visual (V), visuo-movement (VM), and movement (M) related signals, but the nature of information transmitted from FEF and its target within SC remains controversial. The three main models supported by prior studies are illustrated. **(B)** Memory-guided saccade task: a target is briefly presented for 100ms, then after a variable delay of 800-1200ms the fixation spot disappears, instructing the monkey to make a saccade to the remembered target location. **(C)** Time course of the memory-guided saccade task. **(D)** Representative responses of three example SC-projecting neurons in the FEF during the memory-guided saccade task: a Visual neuron (top), a Visuo-movement neuron (middle), and a Movement neuron (bottom). **(E)** Population SDF for SC-projecting FEF neurons categorized as Visual (top, N = 42), Visuo-movement (middle, N = 49), and Movement (bottom, N = 22). For each neuron, the raw SDF was normalized by z-scoring, whereby the mean firing rate was subtracted from each time point and the result was divided by the standard deviation. **(F)** Proportion of SC-projecting FEF neurons out of all FEF neurons categorized using the memory-guided saccade task. The proportions across functional subtypes were not significantly (n.s.) different. **(G)** An example session of laminar recordings in the SC during optical stimulation of the FEF, shown as a heatmap (left). Neurons having excitatory responses with latencies <10ms (green traces at right) were considered monosynaptically activated by FEF stimulation, i.e. FEF-recipient. **(H)** Activity patterns of the same SC neurons from panel g during the memory saccade task, aligned on target onset (left) and saccade onset (right). Green traces are the putative FEF-recipient neurons. **(I)** Proportion of FEF-recipient SC neurons across the three functional subtypes in the SC. ***, p < 0.001.

We used a memory-guided saccade task to dissociate the visual- and motor-related activity^5^ (Figure 3B,C) of neurons. “Visual neurons” responded exclusively following target onset, without activation preceding the saccade. “Visuo-movement neurons” showed both visual responses after target onset and motor-related activity prior to saccade initiation. “Movement neurons” exhibited activation only before saccade onset, with no responses to target presentations. Examples of SC-projecting FEF neurons showing these three activity subtypes are shown in Figure 3D, and pooled data for all such neurons in Figure 3E. In the entire sample of FEF neurons we recorded during the memory-guided saccade task, SC-projecting neurons were distributed uniformly across the three functional subtypes (Figure 3F). They comprised 23.73% of all FEF Visual neurons (42/177), 21.59% of all FEF Visuo-movement neurons (49/227), and 21.78% of all FEF Movement neurons (22/101; p = 0.86). This lack of bias was unchanged when applying a shorter latency-cutoff of <5ms for defining SC-projecting FEF neurons (Figure S3G right). Hence, we found no enrichment of movement-related signal in the FEF output, inconsistent with the motor dominance model (Figure 3A left).

### Signal content of SC neurons that receive FEF input

To discriminate between the visuomotor and no convergence models (Figure 3A, middle vs. right), we performed laminar recordings in the SC while optically stimulating the FEF (Figure 3G). Out of 163 SC neurons recorded across 10 sessions, FEF stimulation modulated a substantial number (N = 64, 39.26%), with excitation dominant (81%, 52/64; Figure S4A). To isolate putative monosynaptic FEF targets, we focused on SC neurons displaying activation <10ms (Figure S4B). They were primarily recorded from the middle channels of the probe (Figure 3G), consistent with the SC intermediate layers. Studying the recorded SC neurons with memory-guided saccades (example raw data across channels in Figure 3H), we found that FEF-driven SC neurons were distributed unequally across the functional categories defined by that task (Figure 3I). They comprised 42.3% (22/52) of the Visuo-movement neurons and 33.3% (15/45) of the Movement neurons we found in the SC, but only 9.6% (5/52) of the Visual neurons (p = 6.86e-04). This functional bias could not be explained by sampling artifacts, because our recordings sampled comparable numbers of Visual (N = 52), Visuo-movement (N = 52) and Movement neurons (N = 45), nor could it be attributed to injection variability, as optical stimulation within the SC to directly identify ChR2-expressing neurons yielded a balanced distribution of all three subtypes (Figure S1D). Hence our results support the visuomotor convergence model: the FEF output to SC conveys a full spectrum of visual-to-motor signals but preferentially targets Visuo-movement and Movement neurons within the SC (Figure 3C middle).

## Discussion

Our findings help to reconcile previous conflicting data from electrical antidromic stimulation approaches^4,6^ by establishing that the primate FEF-SC projection transmits a continuous spectrum of visual-to-motor signals. The present evidence for that conclusion, and against a motor-dominant view of FEF output^4^, was strengthened by our ability to record SC-projecting FEF neurons and other FEF neurons simultaneously to attain unbiased proportions of functional neural types. In the SC, FEF inputs converged primarily onto motor-related neurons, consistent with earlier anterograde stimulation studies^22^. Crucially, retrograde optogenetics offers superior pathway specificity: unlike antidromic stimulation, which may nonspecifically activate nearby axons of unrelated pathways^23^, retrograde viral vectors ensure only the intended projection is labeled and manipulated^11^.

To classify SC-projecting FEF neurons and FEF-recipient SC neurons, we used an activation-latency cutoff of <10 ms, a conservative choice since similar results were obtained with a 5 ms cutoff. Because there is no definitive latency cutoff, some false positives or negatives are theoretically possible. The same is true for FEF-recipient SC neurons. Such issues also affect antidromic electrical stimulation studies; lack of activation does not guarantee lack of projection. Our distributions of neural subtypes, however, were so uniform for FEF (Fig. 3F) and highly skewed for SC (Fig. 3I) that it is unlikely that potential misclassifications could have affected our conclusions.

Overall, these results support a visuomotor convergence model in which diverse FEF outputs preferentially innervate motor-related neurons in the SC. This conclusion favors a view of sensorimotor processing as a continuous, multistage transformation, where activity at any stage along the pathway can advance the computation toward action^24^. Thus, the present results are also consistent with cognitive models proposing graded transitions between perception and action^25,26.^ Future work will be needed to determine whether this organizational principle—convergence of diverse cortical output signals onto motor representations in subcortical structures—can be generalized across cortico-subcortical circuits in other sensorimotor systems.

Our approach was also highly effective at manipulating saccadic behavior. While prior optogenetic studies in monkeys established causal links between neural activity and behaviors, they largely relied on local stimulation at injection sites^27^ or anterograde manipulation^28,29^. Our study fills a critical gap by achieving robust and selective manipulation of a distinct, complete projection. This approach yielded behavioral effects comparable in strength to classical methods including electrical stimulation, but with far greater specificity. These causal results provide a foundation for using the same methods to selectively manipulate circuits for higher cognitive functions^30^.

The vector we used, AAV2-retro, is useful for labeling projection neurons in many areas of primate cerebral cortex^10,31.^ Such neurons are predominantly glutamatergic, and further studies are needed to determine if the vector is as useful for the GABAergic projection neurons that are common subcortically^18,32^,33. A major benefit of retrograde optogenetics is the integrated set of results it provides. Instead of using antidromic stimulation to identify neurons at a projection origin^5,24^,34, orthodromic stimulation to identify neurons at a projection target^21,22,^ and other methods to study the impact of the projection on behavior^29,35,^ all of these findings can be collected in the same animals, with the same methods, at the same time. Applied to primates, as done here, the approach seems promising for circuit-level dissection of higher cognitive functions homologous to those in humans.

## Methods

### Subjects

Two adult male rhesus monkeys (*Macaca mulatta*, monkeys R and A), weighing 9-12kg, participated in the present study. All experimental protocols and procedures were approved by the National Eye Institute Animal Care and Use Committee and complied with the United States Public Health Service Policy on the Humane Care and Use of Laboratory Animals.

### Surgery

We implanted a plastic head holder and two recording chambers for accessing the SC and FEF in the skull under general anesthesia and sterile surgical conditions. The SC recording chamber was positioned on the midline and angled 38 degrees posterior to vertical on the sagittal plane, allowing electrode penetrations to approach the SC surface approximately orthogonally in both monkeys. The FEF recording chamber was placed over the frontal lobe and tilted laterally by 20 degrees. The head holder and recording chambers were secured with ceramic screws and embedded in dental acrylic resin, which covered the top of the skull. After confirming the position of recording chambers using MRI (4.7T, Bruker), a craniotomy was performed for each chamber during another surgery for future recordings.

### Behavioral tasks

All behavioral task events and data acquisition were conducted using custom-made C++ based software (Blip, http://www.robilis.com/blip/). The monkeys were seated in a primate chair with their heads restrained, facing a fronto-parallel screen in a sound-attenuated and electrically shielded room. Visual stimuli were rear projected onto the screen using a digital light processing projector (PJD8353s, ViewSonic). Eye position was monitored at a sampling rate of 1KHz using an infrared eye-tracking system (EyeLink 1000 Plus, SR Research). Water rewards were delivered through a computer-controlled electromagnetic solenoid valve (Parker-Hannifin). The timing of all task-related visual events was recorded relative to the timing of the evoked signal using a photodiode positioned at the corner of the screen.

### Behavior tasks for stimulation

We trained monkeys to perform four different tasks to evaluate the effect of optical or electrical stimulation.

#### Movie free viewing task

The subjects freely watched randomly selected natural history movies (15-25 minutes long) continuously with occasional random water rewards to keep them engaged. Unbeknownst to the subject, the entire viewing period was artificially divided into trials 2-3 seconds long separated by inter-trial intervals of 1-2 seconds. Optical (4.5mW, equals 50mW/mm^2^, 100ms) or electrical (50uA, 100Hz, 100ms) stimulation was applied in half of the total trials, at a random time point between 500ms and 1s after the start of the trials. To ensure wakefulness and engagement, the eye movements were monitored, and the subjects were required to keep their eyes open and within the designated 40° × 40° window throughout the trial. Optical or electrical stimulation was applied only when the eyes were detected within this window. If the animals moved their eyes outside the window or closed them, the current trial was aborted and repeated later. No movement constraints were imposed during the inter-trial intervals. If a subject successfully completed a trial, a small amount of water reward was delivered at the end of the trial. A typical session consisted of 80 or 160 successful trials.

#### Fixation Task

In the fixation task, subjects were required to maintain fixation throughout each trial. At the beginning of a trial, a central fixation point (1° × 1° degree) was illuminated, and the subjects had to fixate on the point within 500ms. Between 500ms and 1000ms after fixation onset, the optical stimulation was applied for 100ms in half of the trials (4.5mW, equivalent to 50mW/mm^2^, 100ms; stimulated trials), while no stimulation was applied in the other half (control trials). Stimulated and control trials were randomly interleaved. Following the stimulation period, subjects continued fixating for an additional 500ms to 1000ms, making the total fixation duration for each trial 2s. The fixation point remained visible throughout the entire fixation period. The fixation window was ±5°. Subjects were allowed to break fixation during the stimulation period (100ms) and 200ms after the stimulated period, as well as during the corresponding 200ms period in control trials. If fixation was broken during this permissive window, subjects had 500ms to re-fixate on the fixation point and complete the remaining fixation period to successfully finish the trial. Successfully maintaining fixation for the required duration resulted in a water reward. A typical session consisted of 80 or 160 trials.

#### Gap Fixation task

The procedure for the gap fixation task was nearly identical to that of the fixation task, except that the fixation point disappeared for 500ms in the middle of the trial. During training, subjects were required to maintain fixation in each trial even when the fixation point was absent. In the stimulation experiment, optical stimulation was applied 200ms after the fixation disappeared and ended 200ms before the end of the fixation-off period. Eye movements during the stimulated period were compared with those during the corresponding fixation-off period in control trials.

#### Delayed saccade task

In the delayed saccade task, subjects were required to make a saccadic eye movement to a visual target that appeared in the periphery after a random delay. Each trial began with the presentation of a central fixation point. After subjects maintained fixation for 500ms to 1000ms, a saccade target was displayed either contralateral or ipsilateral to the stimulated hemifield. The target location was aligned with the saccade vectors elicited by stimulating during the free-viewing task. Following a delay period of 800ms to 1200ms, the fixation point was turned off, serving as the go signal instructing the subject to make a saccadic eye movement toward the target location to receive a liquid reward. Optical stimulation was applied using blue laser light for 200ms, starting with the go signal. Trials with and without stimulation were randomly interleaved. To ensure subthreshold stimulation, the laser power was reduced to less than half (<2mW, equivalent to 20mW/mm^2^). An initial set of 200 trials was first conducted before the formal experimental session, to confirm that the stimulation did not directly trigger saccades (i.e. fewer than 1% of cases, i.e. fewer than one saccade per 100 trials). A typical recording session consisted of 160 - 320 trials in total.

### Behavior tasks for neural recordings

Once units were identified, we first manually mapped their receptive field approximately. We then fixed the target presentation eccentricity and varied the target presentation angle to determine the preferred vector using a receptive field mapping task. Following this, we employed a memory saccade task to classify the functional subtype of each unit.

#### Receptive field mapping task

The procedure of the receptive field mapping task was identical to that of the delayed saccade task except that no stimulation was applied and the target appeared at one of 8 possible locations: 0 degree (up), 45 (left - up), 90 degree (left), 135 degree (left - down), 180 degree (down), 225 degree (right - down), 270 degree (right), or 315 degree (right - up).

#### Memory-guided saccade task

The task began with the appearance of a fixation point. The subject was instructed to acquire and maintain fixation within a 5°×5° window. After holding fixation for 800ms, a target was briefly flashed for 200ms, then the fixation point remained for an additional 800ms –1200ms before disappearing. The offset of the fixation point served as the go signal, prompting the subject to saccade to the remembered target location within 500ms. To successfully trigger the reappearance of the target stimulus, the saccade had to land within 5 degrees of the target location. The subject then had to maintain fixation on the target for another 400ms to receive a reward.

### Viral vector preparation

Adeno-associated viral vectors with retrograde infection expressing the Channelrhodopsin gene (AAV2-Retro hSyn-ChR2-EYFP) were produced by the helper-free triple transfection procedure and purified by affinity chromatography (GE Healthcare). Viral titer was determined by quantitative PCR using Taq-Man technology (Life Technologies). The transfer plasmid (pAAV-hSyn-ChR2-EYFP-WPRE) was constructed by replacing the CMV promoter of the pAAV-CMV-ChR2-EYFP-WPRE plasmid^29^ with the human Synapsin (hSyn) promoter.

### Viral vector injection

We first identified the SC using a combination of magnetic resonance imaging and electrophysiological properties. After the identification, AAV2retro-hSyn-ChR2-EYFP (2.5 x 10^13^ genome copies/mL) was injected into the SC of one hemisphere in both monkeys (monkey R: left SC, monkey A: right SC) using a custom-made injectrode. For each monkey, two penetrations were made at least 1.4mm apart. In each penetration, 1.0ul of the vector was injected at two different depths, separated by at least 500um. The injection process began with an initial infusion of 0.2uL at a rate of 0.4uL/min, followed by a slower infusion rate 0.02uL/min for the remaining volume. This process was controlled using a 10uL Hamilton syringe and a motorized infusion pump (Harvard Apparatus, Holliston, MA, USA). After each injection, the injectrode was left at the site for 1 hour before proceeding to the next site to ensure absorption of viral solution into the surrounding neural tissue. Recording and testing experiments were conducted 2 - 3 month after injection.

### Laser light stimulation

We used custom-made single channel or multi-channel optrodes for optical stimulation and electrophysiology recordings. The single channel optrode had an optic fiber (Doric Lenses, 200um diameter, 0.5NA) attached to a tungsten recording electrode (FHC, 0.5-2MOhm impedance, 125um diameter). The tip of the electrode typically extended 200um beyond the fiber tip. The multi-channel optrodes had the same optic fiber attached to a 24-channel laminar probe (V probe, Plexon INC,100um inter-contact-distance). The tip of the fiber was placed close to the third and fourth channels on the probe. The light source was a 473nm blue diode laser with a maximum power up to 300mW (Topica photonics, iBeam smart). Before the optrode was inserted into the brain, we measured the light intensity at the tip of the optrode using an optical power meter (Newport Corporation, Centauri). During the movie free viewing task, we typically fixed the output power to be 4.5mW (∼50mW/mm^2^) in FEF and 1.5mW (∼16mW/mm^2^) in SC, for significant behavior and neuronal modulation.

### Electrophysiology

Electrophysiological recordings were performed using either single channel or 24-channel V probe optrodes (Plexon). Electrodes were inserted into the brain through a stainless-steel guide tube and advanced using oil-driven micromanipulator (MO-97A, Narishige). Recording sites were targeted using a grid system that enable electrode placement at 1-mm intervals in x and y directions, orthogonal to the guide tube. Extracellular signals were amplified and band-pass-filtered (100 Hz to 8kHz) using a multichannel processor (Plexon). The FEF was identified in the anterior bank of the arcuate sulcus, where saccades could be evoked via low threshold electrical stimulation (Threshold < 50uA, 100Hz) and neurons showed clear visual and/or saccade related responses. The SC was identified as showing clear visual and/or motor responses and clear transition from visual neurons to visual-movement neurons and to movement units during the penetration.

### General experimental procedures

During a typical recording session, once neurons were stabilized and well-isolated, the receptive field of a randomly selected neuron was initially mapped by manually placing a square target at various screen locations to identify the position that elicited the strongest neural responses. At this stage, the receptive field locations were roughly determined by subjectively evaluating the strength of the neural responses through auditory monitoring, and those responses were confirmed as either visual or saccade related. Next, the movie free viewing task was administered to assess the optical modulation of the recorded neurons. The isolated units were then tested on a subset or all of the following tasks: fixation stimulation task, gap fixation task, delayed saccade task, receptive mapping task, and the memory saccade task, among others. Neurons were recorded across all subsequent tasks regardless of whether they exhibited optical modulation, in order to avoid sampling bias.

### Dataset

In total, we recorded 716 neurons from the FEF, comprising 562 neurons from Monkey R across 82 sessions and 154 neurons from Monkey A across 28 sessions, to evaluate the effects of optical modulation during the movie free-viewing task. Of these, 612 neurons were also recorded during the memory saccade task for functional subtype characterization. We excluded 30 neurons due to a mean firing rate of less than 1 spike/s during the task period, and 77 neurons due to the absence of significant visual or saccade-related responses. This resulted in 505 neurons being included for further functional subtype characterization. To assess optical modulation effects in the SC, we recorded 368 neurons (285 from Monkey R in 37 sessions and 83 from Monkey A in 8 sessions). Among these, 323 neurons were also recorded during the memory saccade task. We excluded 23 neurons for low firing rate (< 1 spike/s) and 59 neurons for lacking significant visual or saccade related activity, resulting in 241 SC neurons retained for functional subtype characterization. In addition, we recorded 163 extra SC neurons in 10 sessions while conducting optical stimulation in the FEF, of which 149 were also recorded during the memory saccade task and used for functional subtype characterization.

### Data analysis

#### Spike analysis

Spike sorting was conducted offline using Kilosort^36^ and manually curated by a human expert using Phy2 (www.github.com/cortex-lab/phy) to ensure that the sorted units exhibited physiologically plausible inter-spike interval distributions and waveform shapes consistent with action potentials. Neurons were excluded from analysis if they had low signal-to-noise ratio, a low average trial firing rate (<1spikes/s), or lacked a clear visual or motor responses. The discrete spike times were binned in 10-ms windows and convolved with a Gaussian kernel (σ = 10 ms) to generate the spike density function (SDF) for each unit. In the movie free viewing task, to illustrate the effects of optical stimulation in population, each neuron’s SDF was normalized by dividing by its peak firing rate and then subtracting the average baseline activity (-100ms to 0ms relative to laser onset) before averaging across neurons. In the memory saccade task, SDFs were normalized by z-scoring, whereby the mean firing rate was subtracted from each time point and the result was divided by the standard deviation.

#### Saccade analysis

In the free-viewing task, eye traces spanning 300ms—comprising 100ms before optical stimulation onset, the 100ms optical stimulation period or un-stimulated control period, and the 100ms after stimulation offset—were extracted for saccade analysis. Raw eye position traces were differentiated to obtain velocity and acceleration profile. Saccade detection was based on a modified version of the algorithm described previously^37^. Briefly, candidate saccades were identified when eye velocity exceeded 200 degrees/s and acceleration exceeded 1000 degree/s^2^. Saccades were then detected as the time point when velocity surpassed a threshold of 40 degrees/s. Saccade onset was defined as the time point when velocity first exceeded 40 degrees/s and offset was defined as the time point when velocity fell below this threshold. Sequential saccades were considered valid only if the inter-saccade interval was at least 20ms. For each detected saccade, direction was calculated as the vector from the saccade start to end point, and amplitude was defined as the Euclidean distance between the saccade start to the end point. To minimize noise, only saccades with amplitudes between 5 to 40 degrees were included in the analysis for this task. In the delayed saccade task, reaction time was defined as the interval between the offset of the fixation point and the onset of the saccade towards the visual target.

#### Optical modulation analysis

For optical stimulation experiments, the neural responses of FEF during the 100ms stimulation period were compared to the corresponding 100ms period in the non-stimulated (control) trials. Neurons were labeled as optically modulated if their average responses during stimulation differed significantly from those in control trials, tested via a two-sample t-test using a criterion of p<.05. Neurons showing higher activity during stimulation compared to control were classified as excitatory modulated, while those with lower activity were classified as inhibitory modulated. For significantly modulated neurons, we then measured the modulation latency. Raw spike rasters were divided into two segments: pre-stimulation and stimulation (from onset to offset). Within the stimulation segment, we compared the spike counts in each 10ms bin to the 10ms bin immediately preceding stimulation onset using a paired t-test. The 10ms window was advanced in 1ms steps for successive comparisons. If 5 consecutive windows showed significant differences, the latency was defined as the onset time of the first significant window. The latency was marked as NaN and excluded from the significantly modulated group if no significant window was detected by the end of the stimulation period. Based on the latency, the excitatory modulated population was further divided into 2 groups: (1) fast excitatory neurons with latency <10ms; (2) slow excitatory neurons with latency >=10ms.

#### Functional subtype characterization

We used spike rates from three time windows in the memory saccade task to categorize neurons in the FEF, following the criteria defined previously^38^: a baseline window (−100ms to -50ms relative to target onset), a visual response window (+60ms to +100ms after target onset), and a pre-saccadic window (−100ms from saccade onset up to saccade offset). For each neuron, a one-way Kruskal-Wallis nonparametric ANOVA was performed across trials using spike rates from the three time windows. Post hoc comparisons were then used to assess whether the visual and/or saccadic activity differed significantly from baseline. Neurons showing significant visual but not saccadic modulation were labeled visual neurons (N = 177). Neurons with significant activity in both visual and saccade windows were classified as visual-movement neurons (N = 227). Neurons with significant activity only in the saccade window were labeled movement neurons (N = 103). The characterization for SC neurons used the same approach but with a time window adjusted according to the response properties of SC neurons: baseline window: -100 to -50ms relative to target onset; visual response window: +30 to +100ms after target onset, and the pre-saccadic window: -50ms from saccade onset up to saccade offset.

#### Statistical analysis

For all comparisons of mean values, we used two-tailed two-sample t-tests, or two-tailed paired t-tests for paired comparisons, except for Figure S3B. For Figure S3B, a non-parametric (Kruskal-Wallis) test was used instead. To compare proportions (Figure 2F,I, Figure S1D, Figure S3G right), we used Chi-Square tests.

### Histology

After completing all experiments, monkey R was sedated with ketamine HCl and then deeply anesthetized with an overdose of sodium pentobarbital (50 mg/kg, intravenous). Once the animal reached a fully areflexic state, the transcardial perfusion was performed, beginning with 6 liters of 0.1 M phosphate-buffered saline (PBS, pH7.2), followed by 4L of 4% paraformaldehyde in 0.1M PBS (pH7.2). After perfusion, the brain was cut into blocks in the coronal plane including the midbrain regions. These blocks were postfixed in 4% paraformaldehyde overnight at 4°C, then cryoprotected in a 30% sucrose solution in 0.1 m PBS at 4°C. Once fully saturated and sunk, the tissue was frozen and serially sectioned into 50um slices using a freezing stage sliding microtome (American Optical Company). The tissue sections were then stored in PBS at 4°C until processed for immunofluorescence.

Sections were divided into six evenly spaced series. One series was reserved for assessing endogenous fluorescence while the other five were subjected to immunofluorescent amplification. Detailed procedures for immunofluorescent amplification have been previously published^39^. Briefly, serial free-floating sections were rinsed in PBS, permeated with 0.5% Triton-X 100 in PBS, blocked with 1% Bovine Serum Albumin and Triton-X 100 solution, and incubated in Goat Anti-GFP (1:800, Rockland Cat#: 600-101-215) antibody solution at 4°C on a rocker for 48 hours. Next, the tissue was rinsed and incubated in a secondary antibody solution of 0.5% Triton-X 100, 1% Normal Donkey Serum, and Donkey anti-Goat Alexa Fluor 488 (1:200, ThermoFisher Cat#: A-11055) at room temperature. After three hours, the sections were rinsed in PBS then incubated in Hoechst 33342 (1:1000, ThermoFisher Cat#: 62249) for 10 minutes. Finally, the tissue was rinsed in PBS and mounted on 1% gelatinized microscope slides. Mounted tissue was then dried at room temperature overnight, then dehydrated in graded ethanols (70%, 95%, 100%) and cleared with xylene. The tissue was coverslipped using Cytoseal 60. The sections were then digitized using an Axioscan.Z1 (Carl Zeiss) at 5x using an Axiocam 506 and HXP 120V as a light source using the DAPI, Green Fluorescent, and Texas Red filters.

## Data availability

Data and code to analyze the data have been deposited in a Github repository: https://github.com/xuefeiyu2015/RetrogradeOptogenetics.git and is publicly available as of the date of publication. Any additional information required to reanalyze the data reported in this paper is available from the lead contact upon request.

## Acknowledgments

We thank Nick Nichols, Daniel Yochelson, Denise Parker, and Hayden Warnock for technical support and Carlos Mejias-Apont for assistance in histological preparation of the material. We are also grateful to Anthony Movshon and Mike Hasse from NYU for their insightful discussions and valuable suggestions. This research was in part supported by the Intramural Research Program of the National Institutes of Health (NIH), United States (1ZIAEY000415) and by award number R01NS125843 to M.A.S. from the National Institute Of Neurological Disorders And Stroke of the National Institutes of Health,, and by the Japan Society for the Promotion of Science (JSPS) KAKENHI Grant (23H02781 to K.I) The contributions of the NIH authors were made as part of their official duties as NIH federal employees, are in compliance with agency policy requirements, and are considered Works of the United States Government. However, the findings and conclusions presented in this paper are those of the author(s) and do not necessarily reflect the views of the NIH or the U.S. Department of Health and Human Services.

## Author Contributions

X.Y., A.G. and O.H. contributed to the conceptualization and experimental design. X.Y and A.G. performed the experiments and analyzed the data. K.I. and M.T. produced the viral vector. M.B. and G.K conducted the histology and immunofluorescence histochemistry. X.Y. wrote the original draft of the manuscript. A.G., K.I, M.B., H.N, M.A.S, and M.T. contributed to manuscript editing. All authors contributed to data interpretation and reviewed and approved the final manuscript.

## Declaration of interests

The authors declare no competing financial interests.

## Notes

https://github.com/xuefeiyu2015/RetrogradeOptogenetics

https://doi.org/10.6084/m9.figshare.30141898.v1

### Competing Interest Statement

The authors have declared no competing interest.

### Summary of Updates

Polished the discussion part, making it more concise and add a statement about the limitation of the study

