## Supplementary Information for "Retrograde Optogenetics Reveals Functional Convergence within a Corticotectal Pathway of Non-Human Primates"

**Supplementary Figures**

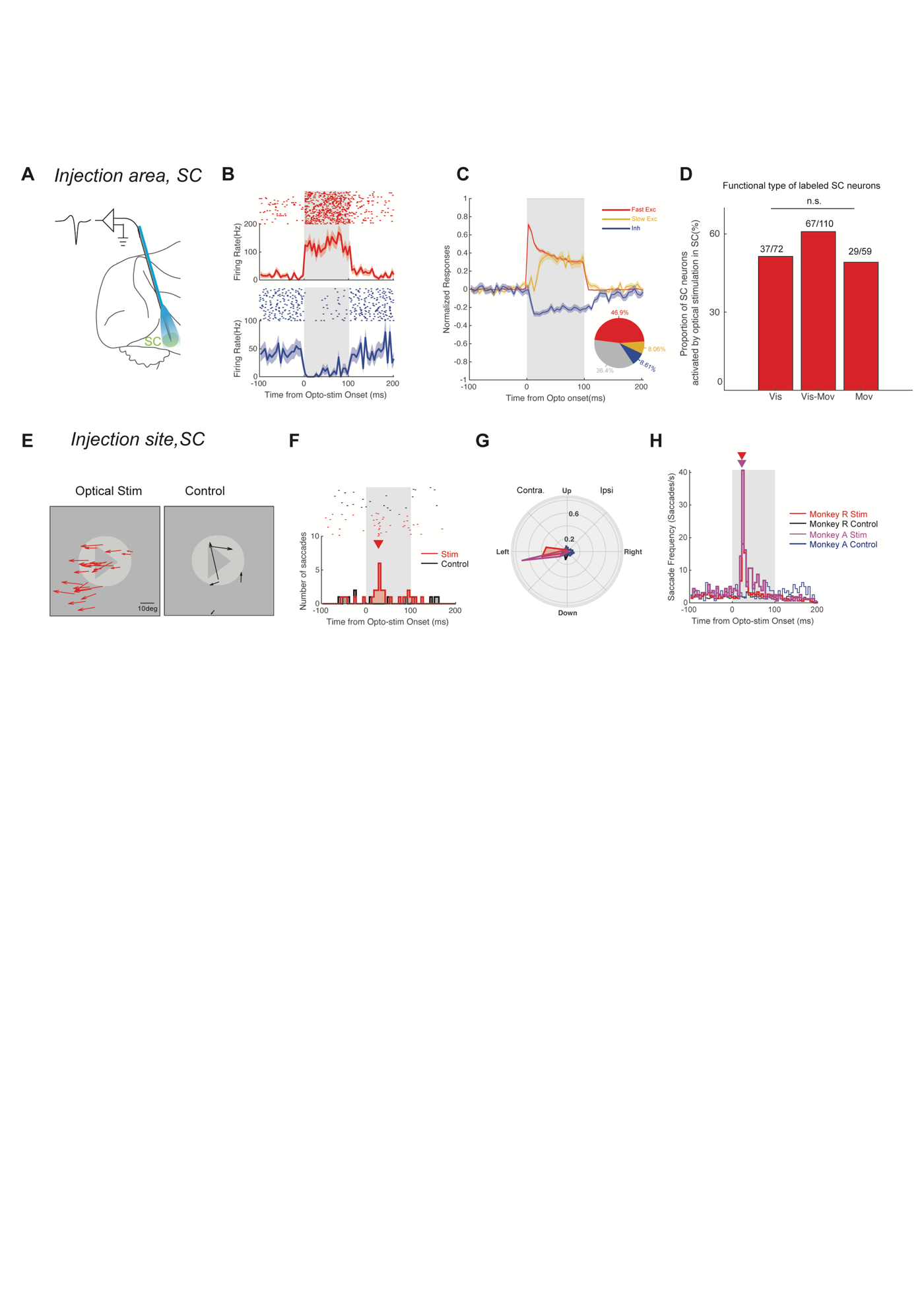

**Figure S1. Optical stimulation in the superior colliculus (SC) induced robust neural and behavior effects. Related to Figure 1 and Figure 2.**

**(A)** A schematic illustrating optical stimulation in the SC. The 473nm blue laser was set with 1.5mW output power resulting in 12mW/mm^2^ at the tip of the fiber. The light power used for SC stimulation was lower than that used for the frontal eye field (FEF) because the SC was the site of viral injection, where expression density was expected higher. Consequently, lower power was sufficient to achieve robust activation.

**(B)** Example SC neurons showing excitation (top) and inhibition (bottom) in response to local optical stimulation.

**(C)** Average population response of optically modulated SC neurons across all recording sessions separated into three groups: Fast Excitation (red), Slow Excitation (yellow), and Inhibition (blue). The pie chart (bottom right) shows the proportion of neurons in each group: Fast Excitation (Fast Exc, red, Latency < 10ms, N = 169); Slow Excitation (Slow Exc, orange, Latency > 10ms, N = 29); Inhibition (Inh, blue, N = 31); and No Effect (gray, N = 131). These results confirm that the optical stimulation effectively modulated SC neurons. The 46.9% of neurons exhibiting fast excitation with short latencies (Latency < 10ms) were likely the directly activated ChR2-labeled SC units. In contrast, neurons showing slower excitation or inhibition likely reflected indirect effects driven by the local circuitry in the SC activated by directly activated ChR2 units.

**(D)** Functional classification of SC neurons directly activated by optical stimulation in the SC. Visual, visual-movement and movement neurons were equally represented among the directly modulated population in the SC. (Visual Neurons: 51.3%, Visuo-Movement Neurons: 60.9%, Movement Neurons: 49.1%; p = 0.25, Chi-Squared Test). This indicates that the ChR2 expression in SC was unbiased across functional subtypes.

**(E)** Example session showing evoked saccade vectors during optical stimulation (left) trials in the SC and control (right) trials.

**(F)** Example session showing peri-stimulus time histogram (PSTH) of saccade frequency (average number of saccades per trial across trials in each 5ms bin) in stimulation (red) and control (black) trials. The triangle indicates the time of peak saccade occurrence (Peak time: 20ms) in this session.

**(G)** Polar histograms showing saccade directions in stimulation and control trials for two monkeys during optical stimulation in the SC. In both monkeys, saccades were directed toward the left visual field, i.e. contralateral to the stimulated SC hemisphere. The histogram is binned in 22.5° intervals. The radial(rho) axis represents the proportion of saccades within each interval relative to the total number saccades recorded for each monkey. When comparing the saccade vectors evoked by SC stimulation to those in Fig.1f, it is apparent that it closely matched those triggered by FEF stimulation. This suggests that the FEF-evoked saccades were mediated through the activation of SC neurons.

**(H)** Population PSTH of saccade occurrence in stimulation and control trials for two monkeys during optical stimulation in the SC. (Peak time: Monkey R: 22.5ms, averaged over 37 sessions and 2183 saccades; Monkey A: 22.5ms, averaged over 8 sessions and 670 saccades.). Saccade frequency was calculated as the average number of saccades within each 5ms bin, divided by the bin width (i.e., saccades per second). Compared to Fig. 1g, the saccade occurrence peak time following SC stimulation occurred earlier than that following FEF stimulation (FEF: 32.5ms), consistent with the SC lying downstream of the FEF in the saccade generation pathway. This latency delay in the FEF is not attributable to differences in light power levels between the SC and FEF, as the power used for FEF stimulation (4.5mW) was much higher than that used for the SC (1.5mW).

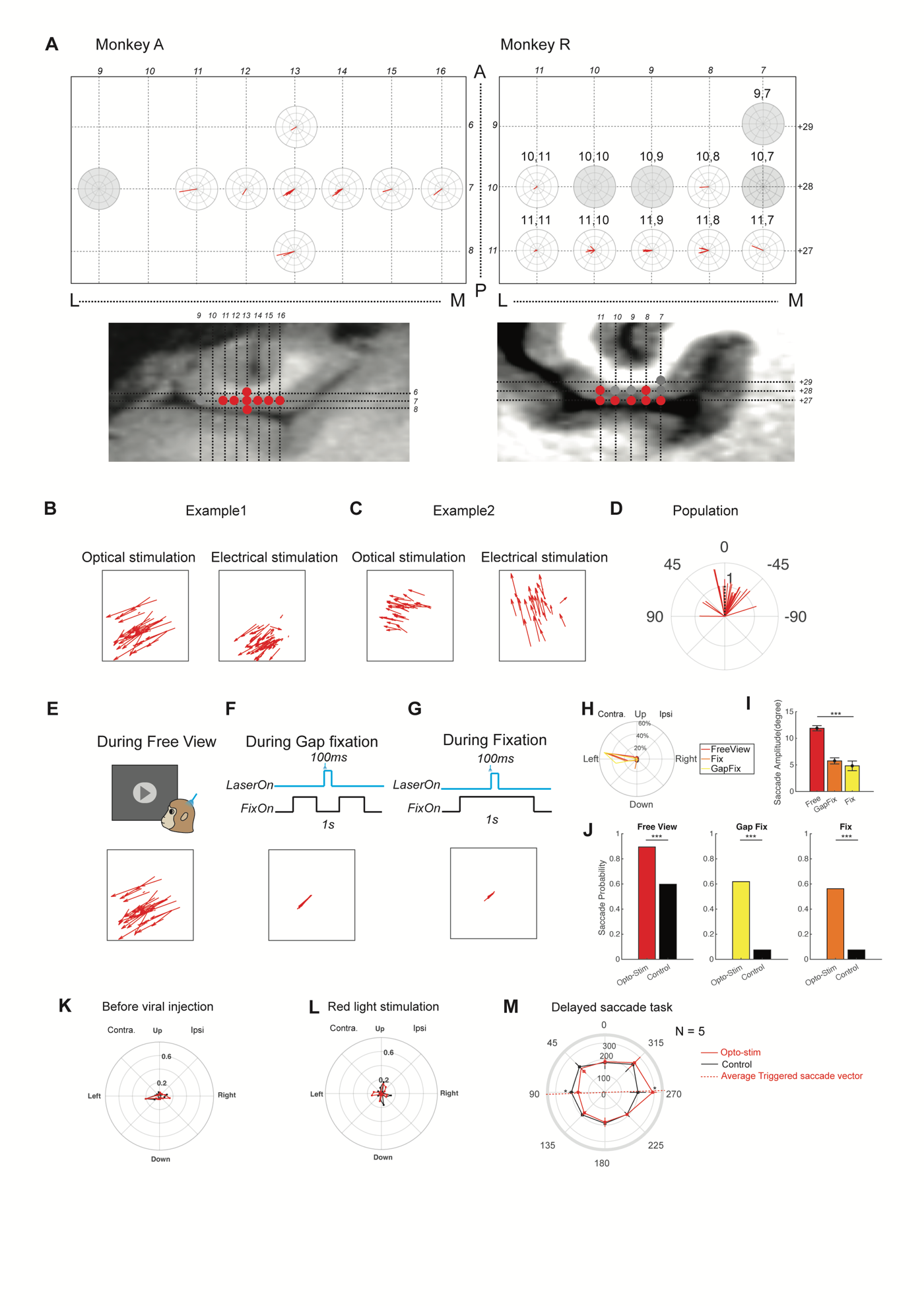

**Figure S2. The FEF optical stimulation reliably evoked saccades across grid locations and task contexts, resembling effects of electrical stimulation, but not under control conditions such as pre-viral injection or red light stimulation. Related to Figure 1.**

**(A)** Saccade vectors evoked by optical stimulation at different grid locations in the FEFs of two monkeys. Each radial distance is an interval of 10 degrees of saccade amplitude, and polar plot directions are relative to the monkey (e.g. left is its left visual field). Left: Monkey A; Right: Monkey R. Images below show the grid locations overlaid on MRIs aligned to the arcuate sulcus (horizontal sections are aligned with the grid plane, 20 degrees tilt from horizontal). Grey polar plots and corresponding dots in the MRI images indicate sites where optical stimulation failed to evoke saccades. In Monkey R, saccade amplitude decreased slightly from medial to lateral sites, while in Monkey A, saccade vectors remained relatively consistent. This variability may be related to the differences in the injection sites and consequent variations in viral expression across animals.

**(B)** An example session in which evoked saccade vectors between optical (left, Average Angle: 126.49° ± 4.50°, [Mean ± SEM], N = 40) and electrical (right, Average Angle: 139.74° ± 6.20°, [Mean ± SEM], N = 52) stimulation was compared to show convergence between two techniques.

**(C)** An example session showing diverging evoked saccade vectors between optical (left, Average Angle: 76.09° ± 1.64°, [Mean ± SEM], N = 31) and electrical (right, Average Angle: 30.25° ± 11.76°, [Mean ± SEM], N = 25) stimulation, illustrating variability across different sites.

**(D)** Comparison of saccade vectors evoked by optical stimulation and electrical stimulation. Polar plots depict the angular differences (theta, Evoked saccade angle (Optical Stim) – Evoked saccade angle (Electrical Stim)) and amplitude ratio (rho, Evoked saccade amplitude (Optical Stim) / Evoked saccade amplitude (Electrical Stim)) of saccades elicited by optical stimulation relative to those evoked by electrical stimulation. Perfect alignment between the two methods would result in an angular difference of 0 degree and an amplitude ratio of 1, as indicated by the black dashed line. Across all cases (N = 26), angular differences remained within 90° (Mean difference: -9.030° ± 6.64°, [Mean ± SEM], comparison to 0: p = 0.80, two-sample t-test) and the evoked saccade amplitudes were similar between these two conditions (Average amplitude ratio: 1.23 ± 0.18; comparison to 1: p = 0.81, two-sample t-test). This slight discrepancy may arise from the differences in the populations of neurons activated by the two methods. During optical stimulation in FEF, only FEF neurons projecting to a particular group of SC neurons were activated, generating saccade vectors consistent with the SC’s coded saccade vector. In contrast, during electrical stimulation in FEF, most neurons distributed near the electrode tip were likely to be activated. This should activate a larger pool of SC neurons, and the final evoked saccade vector would be due to the summation of vectors of all these units. Hence, the optical stimulation is expected to have more spatially restricted effects defined by the projection patterns of optically activated FEF neurons.

**(E-G)** Saccade vectors evoked by optical stimulation in an example FEF site during three behavioral tasks in the same session: e. Free Viewing, f. Gap fixation, and g. Fixation. Across tasks, the evoked saccade directions were consistent.

**(H)** Distribution of saccade angles evoked by optical stimulation in FEF during the three tasks: Free Viewing (red), Fixation (orange) and Gap Fixation Task (yellow). Saccades consistently pointed in the contralateral directions. Number of saccades: Free Viewing: N = 111; Fixation: N = 45; Gap Fixation: N = 99.

**(I)** Average saccade amplitudes during the three tasks: Free Viewing, red, 11.8°±0.49°, Gap Fixation, yellow, 5.75° ±0.58°, Fixation: orange, 4.82°±0.91°. The amplitudes were highest in the free viewing task, followed by the gap fixation task, then fixation task (one-way ANOVA: p = 3.37e-18, F = 47.4, df = 254).

**(J)** Probability of evoking saccades by optical stimulation across different tasks are shown as bar graphs: Free Viewing (left), Gap Fixation (middle), Fixation (right). The saccade probability was computed as the number of trials in which a saccade was generated during the 100ms stimulation period relative to the total number of trials. The probability during control trials was computed within a similar 100ms period. Across tasks, saccade probability was higher in stimulated trials than control trials in Free Viewing (Stim Trials: 89%, Control: 59%, p = 6.9e-8, Chi Squared test), Gap Fixation (Stim Trials: 61.8%, Control: 7.5%, p = 3.68e-11) and Fixation (Stim Trials: 56.2%, Control: 7.5%, p = e-10), reflecting a consistent oculomotor behavior manipulation by optical stimulation under different task contexts.

**(K)** Polar histograms of evoked saccade directions during optical stimulation in FEF before viral injection (470nm, power: 6.5mW, duration: 100ms). The axis is the same as in Fig .1f, i.e. the histogram is binned in 22.5° intervals and the radial (rho) axis represents the proportion of saccades within each interval relative to the total number saccades recorded during the 100ms interval under each condition for each monkey. No direction bias was observed, and the distribution did not differ from control trials. (p = 0.19, two sample Kolmogorov-Smirnov Test, N = 15 sessions).

**(L)** Polar histograms of evoked saccade directions during red-light stimulation (630nm, 4.5mW) in FEF. The axis is the same as in **k**. No direction bias was observed in stimulation or control trials (p = 0.36, two sample Kolmogorov-Smirnov Test, N = 5 sessions). Together with panel **k**, this indicates that the effects seen with blue-light stimulation were not due to thermal or non-specific artifacts.

**(M)** Reaction time differences across 8 directions in a delayed saccade task. The angular coordinate (theta) represents the target angle (°), and the radial coordinate (rho) indicates the reaction time (in ms), averaged across 5 sessions. Supra-threshold optical stimulation at the tested sites evoked similar saccade vectors, the average vector is illustrated by the red dashed line, pointing from 272.4° to 92.4°. When sub-threshold stimulation was applied on the same sites during the delayed saccade task, reaction times were shortest for targets at 90° (left), where the target was positioned aligned to the direction of the evoked saccades. Conversely, reaction times were longest for targets at 270° (right), opposite to the preferred saccade direction. These results suggested that subthreshold stimulation is most effective when aligned with the site’s preferred saccade axis. * p < 0.05, two-sample t-test.

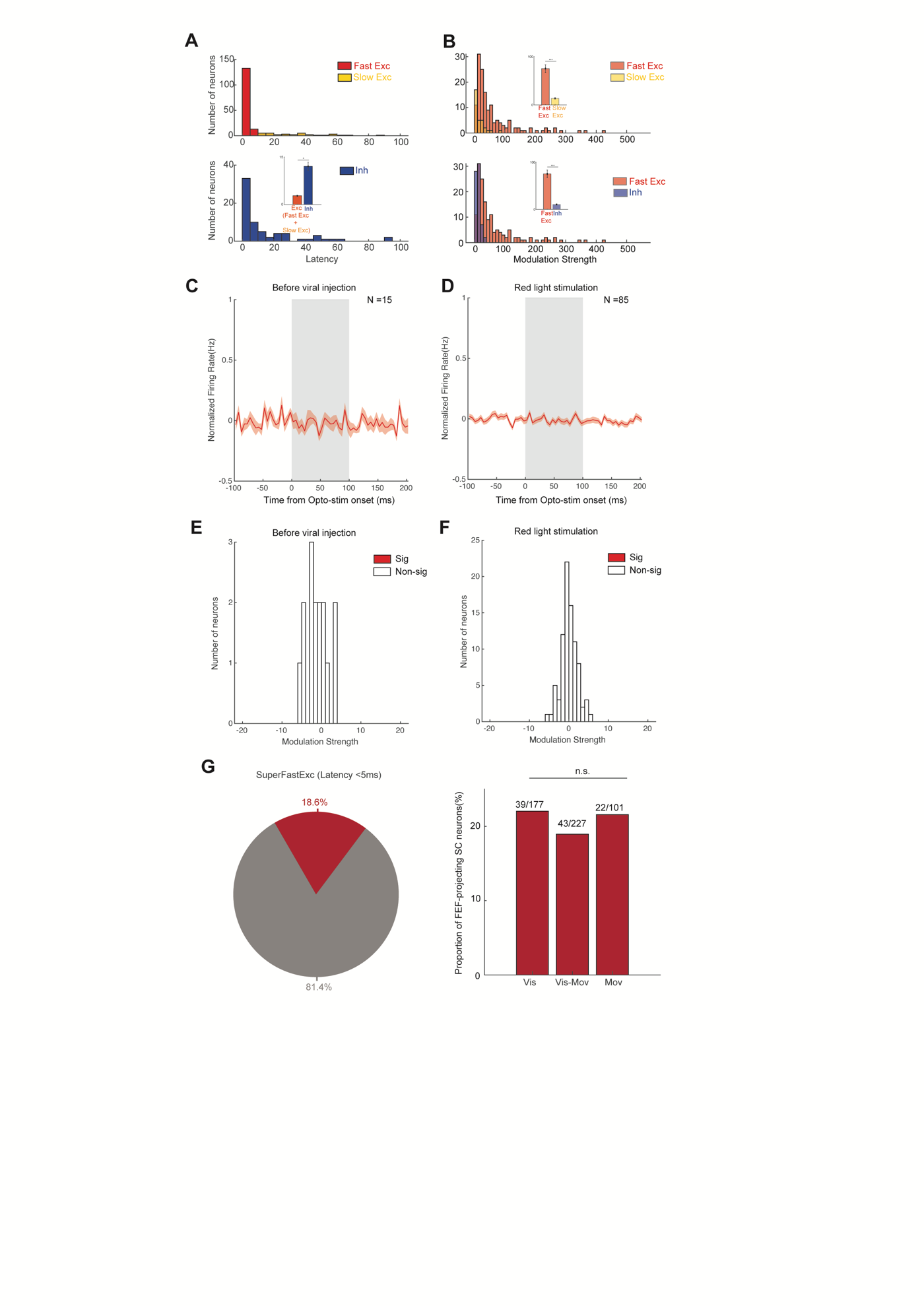

**Figure S3. Additional information on optically modulated sites in the FEF and selectivity of optical stimulation. Related to Figure 2 and Figure 3.**

**(A)** Distribution of modulation latencies among optically modulated FEF neurons. Top: Excitatory units; Bottom: Inhibitory units. Those excitatory neurons that exhibited short-latency responses (< 10ms) likely reflected SC-projecting FEF neurons that were directly activated by optical stimulation (N = 146). In contrast, excitatory neurons with longer latencies (yellow, N =32) and inhibitory neurons (blue, N = 68) were likely modulated via indirect or network-level mechanisms. The latency distributions were significantly different between the excitatory and inhibitory groups (Exc, 6.768 ± 1.14ms; Inhibition, 14.38 ± 2.51ms, p = 0.0018, two sample two tailed t-test). *** p < 0.001.

**(B)** Distribution of modulation strength seen in all optically triggered FEF neurons. The modulation strength was defined as the difference between the average firing rate during the 100ms optical stimulation period and the 100ms pre-stimulation baseline. Top: Comparison of the modulation strength distribution between Fast Excitatory and Slow Excitatory units. Bottom: Comparison of the modulation strength distribution between Fast Excitatory and Inhibitory units. *** p < 0.001 Kruskal-Wallis Test with post-hoc multiple comparisons. Modulation strength: Fast Excitatory (Fast Exc), 65.54 ± 7.14, Slow Excitatory (Slow Exc): 16.32 ± 3.03, Inhibitory (Inh): 12.44 ± 0.87, p =7.84e-36, Kruskal-Wallis Test. Optical stimulation evoked significantly stronger modulation in the Fast Excitatory units compared to both the Slow Excitatory units and Inhibitory units. Combined with the latency differences shown in **(A)**, these results suggest that Fast Excitatory units are directly activated by optical stimulation, whereas Slow Excitatory and Inhibitory responses likely reflect indirect effects, characterized by slower latency and weaker modulation strength.

**(C)** Average PSTH during optical stimulation in free viewing tasks before viral injection (Blue light: 6.5mW, 100ms). The shaded area marks the optical stimulation period. There was no significant modulation observed during this period. N = 15 units.

**(D)** Average PSTH during red light stimulation (630nm, 4.5mW, 100ms), which should not activate ChR2, in a free viewing task conducted a few minutes after the blue light stimulation session. No modulation was observed. (N = 85 units).

**(E)** Distribution of modulation strength across units recorded before viral injection. Across the population, no significant modulation was detected. (-1.21 ± 0.18, [Mean±S.E.M.], compared to zero: p = 0.67, two sample t-test, N = 15)

**(F)** Distribution of modulation strength during red light stimulation. No significant modulation was detected in the population. (0.032 ± 0.024, [Mean ± S.E.M], compared to zero: p = 0.99, two-sample t-test, N = 85). Together with panel **e**, these results confirmed that the observed effects from blue-light stimulation were specific to ChR2 expression and not due to thermal or nonspecific optical artifacts.

**(G)** Proportion of three functional subtypes among SC-projecting FEF neurons (right) defined using a latency threshold of <5ms during optical stimulation (left). Despite this stricter criterion, all three functional subtypes remained broadly represented among the SC-projecting FEF neurons (Visual Neuron: 22.03% (39/177), Visual Movement Neuron: 18.94% (43/227), Movement Neuron:21.57% (22/102), test for distribution difference: p = 0.71, Chi-Squared Test). This suggests that our support for the visuomotor convergence model is robust and not dependent on the specific latency threshold used to define optically labelled SC-projecting neurons.

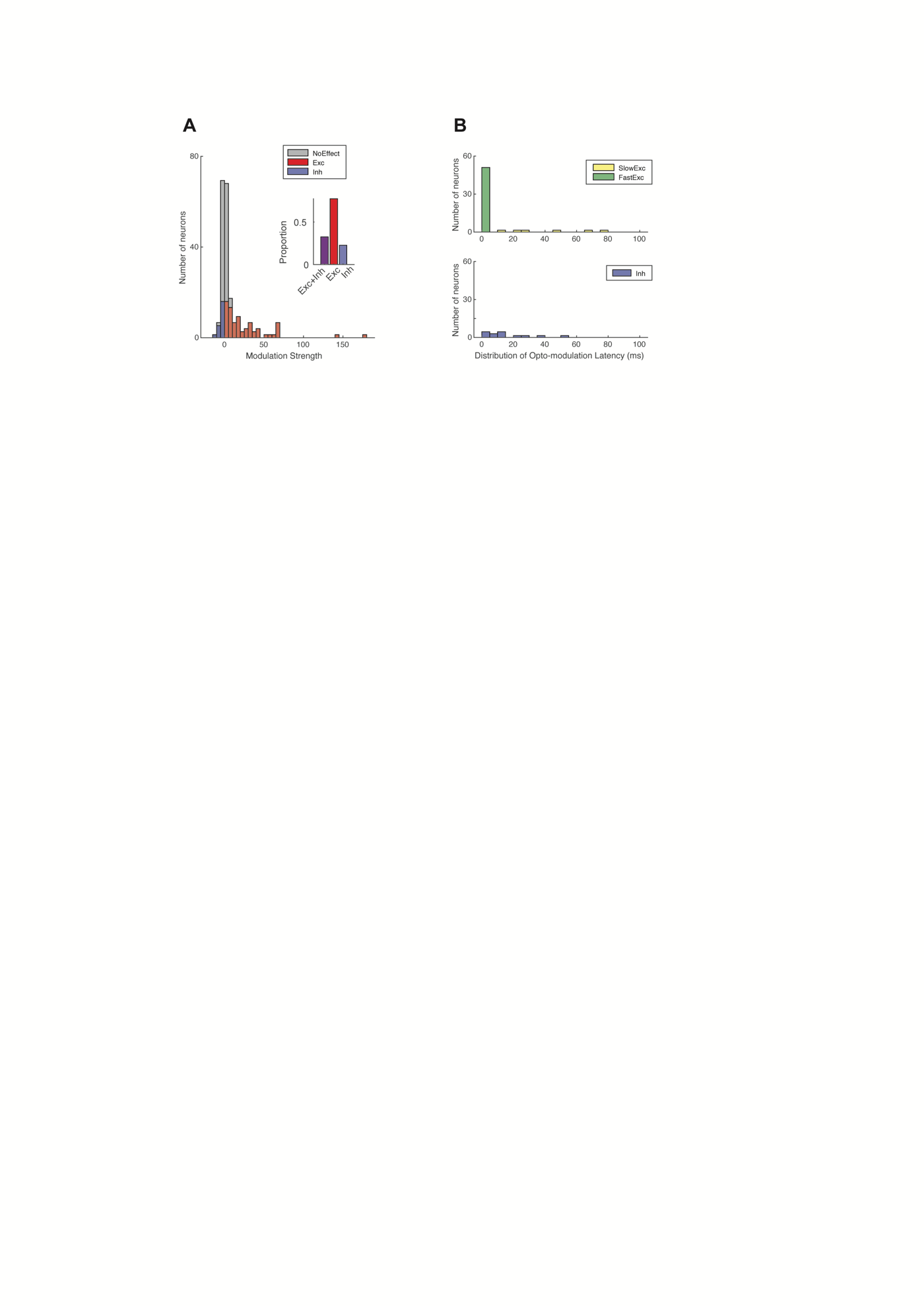

**Figure S4. Modulation of SC neurons during FEF optical stimulation, and control conditions. Related to Figure 3.**

**(A)** Distribution of modulation strength for SC neurons during FEF optical stimulation. Modulation strength is defined as in the Supplementary Fig.3b. Red bars indicate significantly excited units, while blue bars indicate significantly inhibited units. The inset shows the proportion of modulated SC units (39.26%, N = 64 among 163), with 81% of those exhibiting excitatory modulation (Excitation: N = 52; Inhibition: N = 12).

**(B)** Distribution of modulation latency for excitatory (upper panel) and inhibitory (lower panel) SC units. N= 64. Those excitatory units that showed modulation latency within 10ms (N = 42) were defined as SC neurons directly activated via FEF projections.
